# Top-down and bottom-up processes jointly explain mesopredator movement and foraging ecology

**DOI:** 10.1101/2023.04.27.538582

**Authors:** Katie R.N. Florko, Tyler R. Ross, Steven H. Ferguson, Joseph M. Northrup, Martyn E. Obbard, Gregory W. Thiemann, David J. Yurkowski, Marie Auger-Méthé

## Abstract

Prey availability and predation risk drive animal distribution, movement, and foraging ecology, yet studies rarely analyze multiple predator-prey levels together. Understanding how predators optimize risk-reward tradeoffs is important for species conservation and management, especially in systems facing extreme ecosystem change. We examined how top-down (modelled polar bear habitat selection) and bottom-up (modeled fish diversity) processes influence the habitat selection, movement, and foraging behavior of 26 ringed seals (greater than 70,000 dives and 10,000 locations over 877 seal days). Our results suggest that polar bears spatially restrict seal movements and reduce the time seals spend foraging, potentially decreasing foraging success. Seals were more likely to be present and dive longer in high-predation risk areas when prey diversity was high. Further, seal habitat selection models excluding polar bears overestimated core space use. These findings illustrate the dynamic tradeoffs that mesopredators make when balancing predation risk and resource acquisition.

## Introduction

Balancing the demands to forage and avoid predators shapes the spatial distribution, behavior, and habitat use of animals (e.g., 1–3). Optimal foraging theory posits that prey select areas with plentiful resources (4) and avoid regions where predation risk is high (e.g., the landscape of fear, (5)). In some cases, individuals avoid areas of abundant resources due to the risk of predation, producing non-consumptive effects from foregone foraging opportunities (6). In addition to these spatial tradeoffs between predation risk and foraging resources, prey will also allocate time and energy to antipredator behaviors (e.g., vigilance) during – or in lieu of – foraging (2, 7). The influence of predation risk on foraging behaviors can have significant consequences for energetic gain and expenditure (6). Thus, it is critical to incorporate information on both predation risk and prey distribution when aiming to understand animal ecology and the space use and behavioral tradeoffs animals make between food and risk. Given the rapidly changing environmental conditions and increasing anthropogenic pressures on natural ecosystems (e.g., (8)), it is critical to understand predator-prey dynamics and their role in influencing habitat use and movement decisions of vulnerable species (9, 10).

Animal tracking is an essential and increasingly accessible tool to assess movement and behavior, especially for wide-ranging or cryptic species (11–13) and the resulting data can be used to understand habitat selection, movement, and space use (14, 15). While other methods such as giving-up densities and camera traps have provided valuable insight into predator–prey dynamics, particularly in terrestrial systems, tracking data offer individual-level, continuous observations across broad spatial and temporal scales (11). Further, as multi-species tracking studies become more common, these data provide an unprecedented opportunity for simultaneous assessment of top-down and bottom-up influences on species. Such studies, and modelled data, have the potential to resolve longstanding uncertainties in animal ecology, where studies often focus on bottom-up processes only, potentially clouding inference and misrepresenting space use metrics used to inform conservation efforts (e.g., identification of critical habitat).

Polar bears (*Ursus maritimus*) and ringed seals (*Pusa hispida*) form a strong predator-prey relationship (16) where polar bears are the main predator of ringed seals and ringed seals are the main prey of polar bears (e.g., (17)). Spatial variation in perceived risk from polar bears is likely to have strong effects on seal behavior according to the criteria of (18): 1) both species share a heterogeneous physical landscape associated with sea ice, 2) polar bears use an ambush-style of hunting (i.e., “sit and wait”, (19)), and 3) ringed seals lack morphological defenses against predation (20). At the end of a dive, seals must return to the surface to breathe through one of the multiple holes they maintain in the ice during the ice-covered season, providing opportunities for polar bears to capture them (21). Ringed seals maintain haul-out lairs between the sea ice and snow for shelter and breathing, but polar bears commonly capture seals in these lairs (21). Predators have previously been found to affect the diving patterns of ice-associated prey. For example, bowhead whale (*Balaena mysticetus*) dives were deeper and shorter when predation risk from killer whales (*Orcinus orca*) was high (22). Similarly, we expect ringed seals to modify their diving behavior to mitigate the risk of polar bear predation (see below).

Climate change threatens ringed seals and polar bears by reducing the snow and sea ice needed for birth lairs and foraging, while also shifting the prey regime for seals (23–27). Across the Arctic, lower trophic levels are shifting in species composition, quality, and distribution, with changes projected to accelerate (26–29). The southernmost regions of the Arctic are experiencing rapid climate warming and sea ice loss, making them important indicators of ecosystem change. In Hudson Bay, declines in the abundance and distribution of energy-rich Arctic cod (*Boreogadus saida*) and increases in the abundance of smaller temperate-associated fish have been predicted, with an overall increase in total prey biomass and diversity (29). These changes are projected to continue, with important implications for seal foraging success (29). For ringed seals, an annual index of prey diversity – interpreted as an integrative proxy for prey availability across seasons – was the strongest predictor of space use during the ice-free season, when polar bear pressure is low (30). Prey diversity is a valuable metric because, consistent with the portfolio effect, diverse communities stabilize foraging opportunities despite seasonal shifts in individual species, and even when estimated at coarse annual resolutions, they capture integrative spatial patterns across seasons that can be meaningfully matched with finer-resolution predator data (30–32). Simultaneously studying ringed seals, polar bear predation risk during the ice-covered season, and fish abundance and diversity is critical to understanding trophic relationships and for effective conservation (9).

To build on classic risk-reward theory, we examine how prey diversity can alleviate predator-imposed constraints on mesopredator behavior. Specifically, we analyzed satellite telemetry data from 26 ringed seals (over 70,000 dives and 10,000 predicted locations) collected during the ice-covered months over three years (2010-2012), and, as a proxy of predation risk, daily estimates of top predator habitat selection derived from the movement data of 39 polar bears (over 18,000 locations) in Hudson Bay, Canada. We also included modeled annual prey (fish) biomass and diversity (from (29)) and daily sea ice concentration (Fig. 1). Prey diversity has been shown to increase consumption rates and strengthen predator foraging responses, suggesting that diverse communities elevate foraging reward (33, 34).

**Fig. 1.**
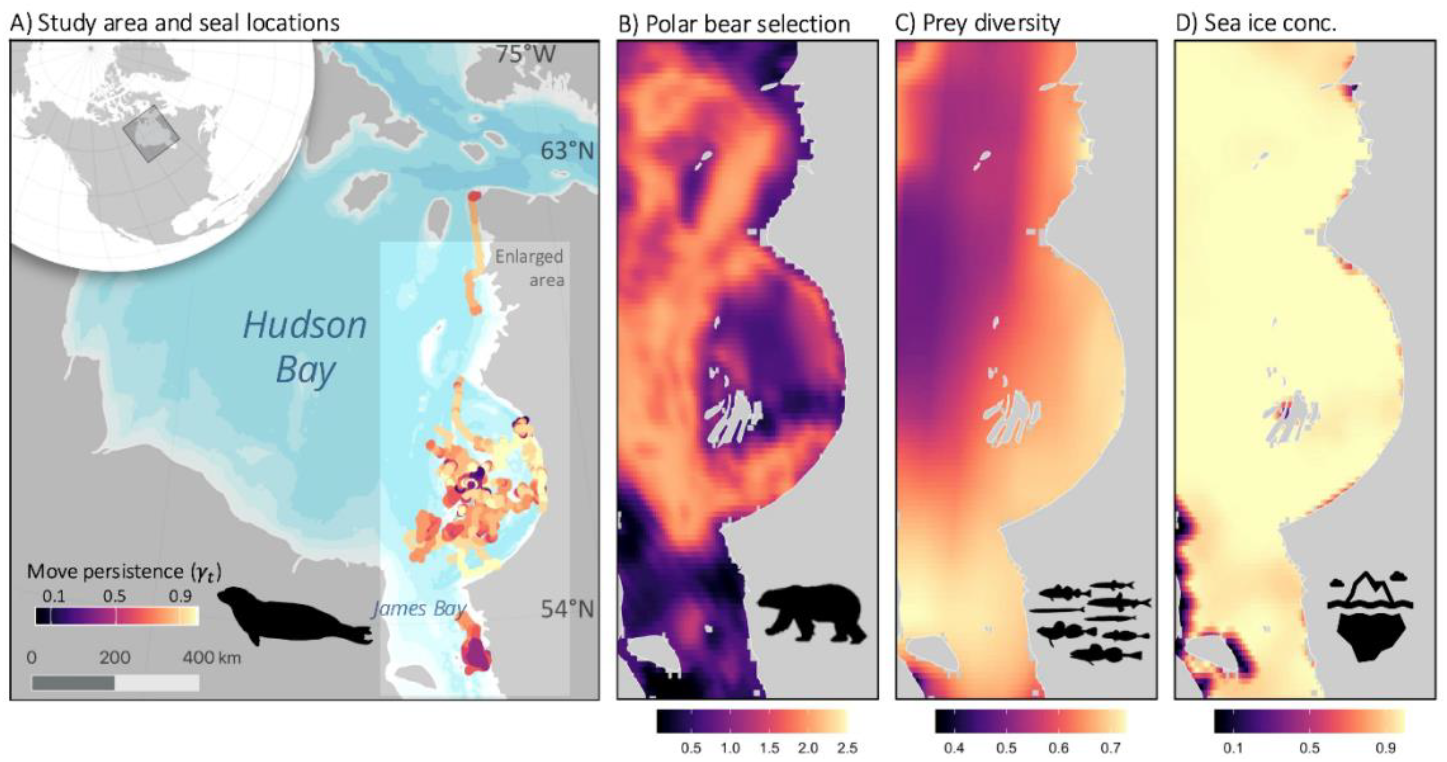
Maps of ringed seal, polar bear, fish diversity, and environmental data. (**A**) Map of the study area with ringed seal predicted locations colored by estimated move persistence. Maps of enlarged study area coloured by (**B**) predicted polar bear selection, (**C**) prey diversity (bilinearly interpolated at the scale of the polar bear selection and sea ice resolution for visualization), and (**D**) sea ice concentration in eastern Hudson Bay, Canada. Panel-specific colour scales are presented at the bottom of each panel. Polar bear selection and sea ice concentration are estimated daily; this is an example day, 15 February 2013. Prey data are estimated yearly; this is prey diversity in 2013. Presented ringed seal movement in (**A**) are for the full study period.

We expected that ringed seals would avoid using riskier areas, and instead use areas with low predation risk and high prey biomass and diversity. Further, we expected that the trade-off between risk and reward would be mediated by prey diversity, such that seals might tolerate high predation risk when prey density or diversity – and therefore foraging reward – is high, consistent with the hazardous duty pay hypothesis (6). Additionally, we hypothesized that when predation risk was high, ringed seals would dive less frequently to reduce the number of ascents to the surface, ascend slower and shorten their dives to allow for flexibility in reaching alternative breathing sites. Alternatively, we hypothesized that ringed seals would lengthen their dives when risk is high to reduce the required number of dives (and their surfacings) to reach stomach capacity (Fig. 2). Our study advances our understanding of predator-mediated foraging ecology in a rapidly-changing ecosystem and highlights the importance of incorporating predator effects when identifying core habitat for conservation.

**Fig. 2.**
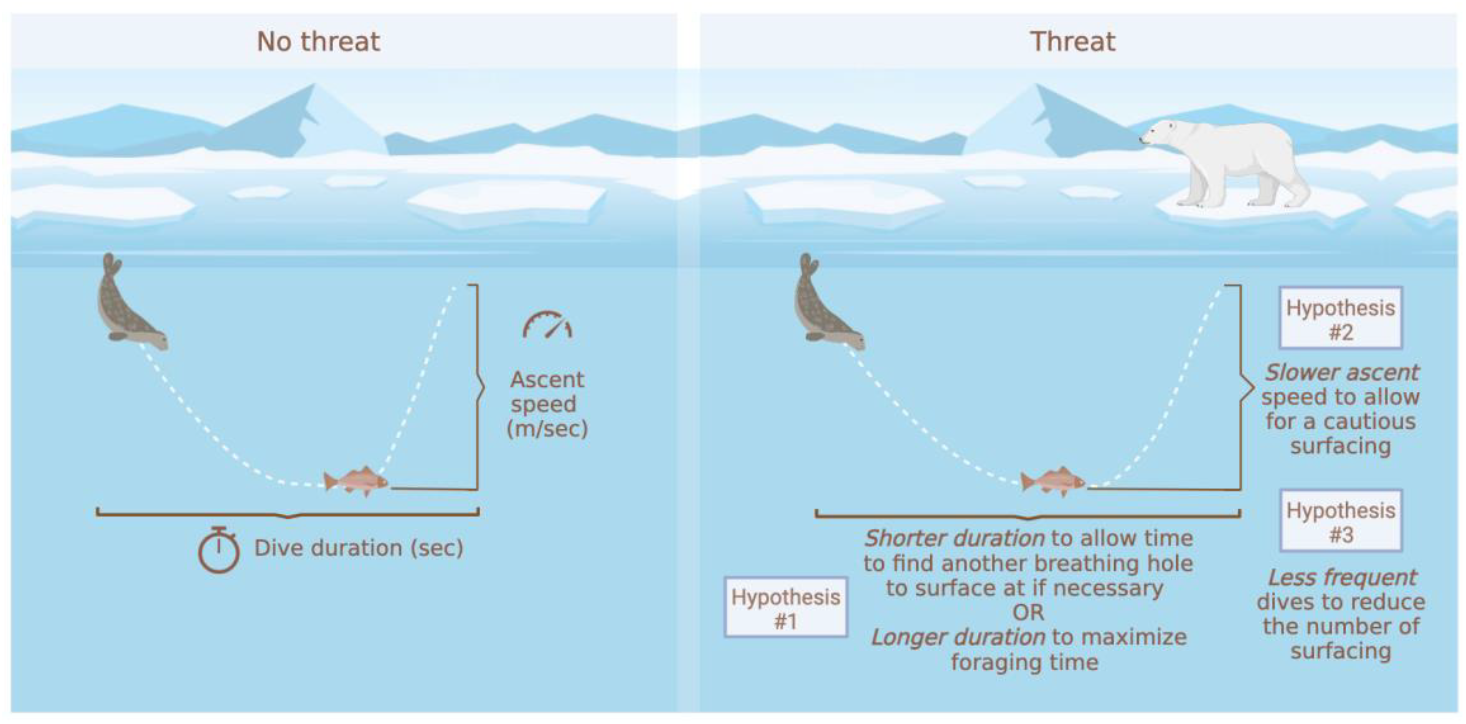
Conceptual diagram of hypotheses. Conceptual representation of three hypotheses for how perceived risk of polar bears at the surface may influence ringed seal diving behavior.

## Materials and Methods

### Seal transmitter deployment

We captured, sampled, and tagged 26 ringed seals in August-October of 2010-2012 using mesh monofilament nets set perpendicular to the shore in shallow water (<8 m depth) in eastern Hudson Bay near the coastline of the Belcher Islands, Nunavut, Canada (Table S1). We equipped ringed seals with Argos satellite telemetry transmitters (9000x data loggers) from the Sea Mammal Research Unit (University of St. Andrews, UK), which recorded location and summary metrics for each dive.

### Seal state-space modeling

Argos locations provided by the tags are associated with error (35, 36), thus we used a state-space model to predict locations that account for this error and used these predicted locations in all subsequent analyses. Specifically, we used the aniMotum R package version 1.1 (36) to fit a continuous-time correlated random walk state-space model that used irregularly sampled locations with error to produce predicted location data at a 2-hr interval. Before applying the model, we split tracks into smaller segments when transmission halted for >12 hrs. The smaller segments were assigned a unique identification number. We removed segments with <100 locations as these led to model convergence issues. Before applying the state-space model, the mean interval between consecutive locations was 1.32 ± 0.24 (median ± standard deviation) hours. Long gaps (>10 hours) were infrequent (0.44% of intervals). The state-space model applied to this data (n = 16,719 raw locations) returned 10,022 predicted locations over a total of 877 seal days during the ice season (i.e., from freeze-up [approx. early December] through break-up [approx. late June]). We used one-step-ahead residuals to assess the goodness-of-fit of our state-space model (36, 37). Additionally, we used the aniMotum::sim_post() function to simulate alternative track locations (n = 100 repetitions) based on the fit state-space model. These simulations allowed us to incorporate location uncertainty into our seal habitat selection models (see below).

#### Seal dive data

The tags also transmitted summarized metrics for each dive deeper than 2 m, including duration, maximum depth (herein “dive depth”), and five intermediate depth points for each dive along with the percentage of the duration elapsed since the beginning of the dive at each intermediate depth point. We calculated the ascent speed (meters per second) as the depth before the ascent divided by the duration from the last depth point to the end of the dive. We calculated dive frequency as the number of dives per seal per day. Each dive also had an associated date and time. We estimated the location of each dive by using the date and time of the dive and linearly interpolating along the state-space model-predicted movement track for each animal. There were 71,942 dives that were matched to the predicted movement locations for use in the dive analysis.

### Covariates for ringed seal habitat use, movement, and dive models

Each seal location (including dives) was spatially and temporally matched to corresponding covariates in downstream analyses, including polar bear selection, prey prediction, and environmental data; covariates detailed below.

#### Predator (polar bear) predictions

We captured and equipped 39 female polar bears with GPS collars in southern Hudson Bay between 2007 to 2013 (n = 18,226 locations). We fit resource selection functions using combinations of sea ice concentration, bathymetry, and distance to coast, where the final model was fit using a Least Absolute Shrinkage and Selection Operator (LASSO) framework (38, 39)(details in Supplementary Material). We predicted daily polar bear habitat selection (herein *polar bear selection*) rasters for each day from the resource selection function (40, 41), and used a spatial bilinear interpolation to match each ringed seal location and dive data with the corresponding value for polar bear selection for the respective day.

#### Prey (fish) predictions

We used annual modeled ringed seal prey biomass data (0.5° resolution) provided by (29, 42) for three prey species that were shown to explain the movement ecology of Hudson Bay ringed seals during the open-water season of the same study years (29, 30): Arctic cod (*Boreogadis saida*), capelin (*Mallotus villosus*), and northern sand lance (*Ammodytes dubius*). In addition, we used Simpson’s Diversity Index (herein *“prey diversity”*) and total biomass of the eight most common ringed seal fish prey species, as in (30). The spatial prey biomass data was modeled using a dynamic bioclimate envelope model (DBEM) that used changes in ocean conditions to calculate annual changes in growth, population dynamics, habitat suitability, and movement for each species (29). Briefly, the DBEM assessed how climate change scenarios impact biogeographic characteristics (43). The DBEM relied on Earth system model data from the Intergovernmental Panel on Climate Change’s (IPCC) Coupled Model Intercomparison Project (CMIP5), specifically oceanographic data, provided by three earth system models (GFDL, IPSL, and MPI), and mechanistically models how changes in oceanographic conditions affect each species’ (separately) physiology, growth, population dynamics, and movement (44, 45). The DBEM predicts yearly abundance and biomass for each species on a 0.5° longitude by 0.5° latitude grid (see (29) and Supplementary Material for further details). We matched each ringed seal location and dive with prey biomass and diversity using a spatial bilinear interpolation for the respective year.

#### Environmental data

We obtained daily sea ice concentration data (25 km resolution) from the National Snow and Ice Data Centre (NSIDC; www.nsidc.org) Nimbus-7 SMMR and DMSP SSM/I-SSMIS Passive Microwave Data satellites. We rasterized and extracted (using a bilinear interpolation) the sea ice concentration value associated with each ringed seal location and dive using the raster package in R (46). We obtained bathymetry data (0.01° resolution) for each ringed seal location and dive from the National Oceanic and Atmospheric Administration (NOAA) Environmental Research Division Data Access Program (ERDDAP) data server *etopo180* (47) using the rerddapXtracto package in R (48).

### Modeling seal habitat selection

We were interested in the potential effects of polar bears on ringed seal habitat selection within their home ranges and thus used a resource selection function (49, 50). Resource selection functions compare locations of an animal to locations deemed to be available for them (51, 52). We addressed the potential issue of pseudoreplication by thinning our dataset to one randomly selected predicted location per seal per day (52). While we focus on the results of this thinned data set, we show that an analysis on the full dataset of predicted ringed seal locations at a 2-hr time-step provides similar results (Tables S7-S8). The available locations were randomly generated locations (n = 25 per used location, (53)) for which we extracted all spatial covariate values. The available locations were sampled from a minimum convex polygon developed using all locations for all seals with a 30-km buffer and clipped to exclude land. We determined the number of available points using a coefficient stabilization analysis, where we fit models with 1-40 (by an interval of 5) available points per used location, and visually inspected at which number of available points the coefficient estimates stabilized (54). The resource selection functions model the binary response (use/available) relative to the explanatory covariates (i.e., combinations of sea ice concentration, polar bear, and prey covariates) using a logistic regression (a generalized linear model with a binomial distribution and logit link) that we fit using the glmmTMB R package (55).

We fit resource selection functions with various combinations of polar bear, sea ice concentration, and prey covariates to explore their ability to explain variation in ringed seal space use at the home-range scale. Our primary objectives were to assess how polar bears may affect ringed seal habitat selection, thus, we used a model selection approach that considered only models that could help address this question and were ecologically relevant. First, since our prey metrics were correlated with each other, we determined which prey metric was most important at the home-range scale by fitting and ranking five models each with one single explanatory variable: biomass of Arctic cod, biomass of northern sand lance, biomass of capelin, total prey biomass, or prey diversity. We used this “most important fish’’ metric to represent bottom-up influence in our subsequent analyses. Next, we fit models with all combinations of three covariates: sea ice concentration, best prey metric, and polar bear; as well as models with an interaction between prey and polar bear covariates (n = 9 models). We also fit a null model for comparison. All models (including the null) included bathymetry as a fixed effect to account for physical constraints in the seal’s diving behavior and thus access to prey. To demonstrate how the inclusion of trophic interactions affects mesopredators space use predictions, we made three prediction maps for ringed seal habitat selection: 1) based on a model that included the best prey metric as a covariate, 2) based on a model that included polar bear as a covariate, and 3) based on a model that included the interaction between the best prey and polar bear covariates.

We checked for correlation between covariates prior to modeling and found no signs of strong correlation (i.e., > 0.6, Table S3). We ranked models using Akaike’s information criterion with small sample correction (AICc), where the model with the lowest AICc value was considered best. However, if another model was within two ΔAICc of the lowest AICc model and had fewer parameters, it was deemed more parsimonious and therefore “better” (56).

To assess the predictive ability of our model, we used the leave-one-individual-out cross validation analysis described by (57) on our best-supported resource selection function model. Briefly, for each individual, we removed its data and used the remaining individuals as a training dataset for our model. We then used the estimated model parameters to predict the resource selection function values at the locations used by the individual that was omitted. We counted the number of predictions that fell in each of ten even-increment bins (“RSF bins”), and calculated the Spearman Rank correlation between the RSF bins against the area-adjusted frequency of used locations for each individual.

To propagate the location error from the ringed seal Argos data in our habitat analysis, we fit our best resource selection function to each repetition (simulated based on the fit state-space model, see above) and created a 95% confidence interval based on the resulting distribution of coefficient estimates.

### Modeling seal movement behavior

To understand the potential effects of polar bears and prey on ringed seal movement behavior, we fit a move-persistence mixed model on predicted ringed seal locations using the mpmm R package (58). The model provides an estimate of directional persistence along the movement tracks and is a continuous value between zero and one where lower values represent low directional persistence, likely associated with residency behaviors (e.g., area-restricted search, foraging, resting), and higher values represent high directional persistence, likely associated with traveling (58, 59). This metric has been widely used and validated in marine mammals to infer behavioral states from movement alone (e.g., (30, 58, 60)), allowing us to capture fine-scale shifts between searching and traveling behaviors. We included the polar bear and best prey covariates separately and together (n = 3 models); our model formulations were limited to first-order terms as the mpmm package does not accommodate interactions. We also included bathymetry as a covariate in all models to account for structural constraints to their movement. We ranked these three models using AICc, as above.

### Modeling seal diving behavior

To understand how prey and polar bears may affect ringed seal diving patterns, we quantified the relationship between three dive metrics, dive frequency, duration, and ascent speed, and prey and polar bear covariates. To do so, we used linear mixed-effects models fit with the glmmTMB R package. To reduce autocorrelation, we selected every 10th dive for each individual. Additionally, each model included a first-order (AR1) autocorrelation structure to further account for the time-series nature of the dive observations. Similar to our resource selection function models, we fit all combinations of sea ice concentration, best prey metric, and polar bear covariates, as well as an interaction between prey and polar bears (n = 9 models per diving behavior). For consistency with the resource selection function methods, we used the same pre-selected prey metric (40). Finally, for each set of models, we fit a null model for comparison. All models, including the null models, included a random intercept term for ringed seal identification number to account for individual variation and dive depth as a fixed effect to account for physiological limitations of diving (61). We ranked models using AICc in the same approach outlined for the resource selection functions.

## Results

### Ringed seal habitat selection, prediction maps, and movement

Our best-supported model describing ringed seal habitat selection (hereafter referred to as *ringed seal selection*) included an interaction between the predicted values from the polar bear resource selection function (hereafter referred to as *polar bear selection*) and prey diversity (Table S4). As polar bear selection increased, ringed seal selection decreased, and for areas with high polar bear selection values, ringed seal selection was more likely if prey diversity was high (Fig. 3A). We also found that ringed seal movement appeared to be constrained to areas with low polar bear selection values (Movie S1). Additionally, this model indicated a positive relationship with bathymetry and sea-ice concentration, with higher ringed seal selection values in shallow and high sea-ice concentration areas; Table S5). This model performed well during our leave-one-out cross validation, where the resource selection function bins were highly correlated with the area-adjusted frequency (0.94; Fig. S2), indicating high model prediction accuracy (57). Importantly, this model, which included top-down (polar bear), bottom-up (prey diversity), structural (bathymetry), and environmental (sea ice concentration) covariates, yields a prediction map that differs from the one predicted when top-down or bottom-up pressures are excluded from the best model (Fig. 4).

**Fig. 3.**
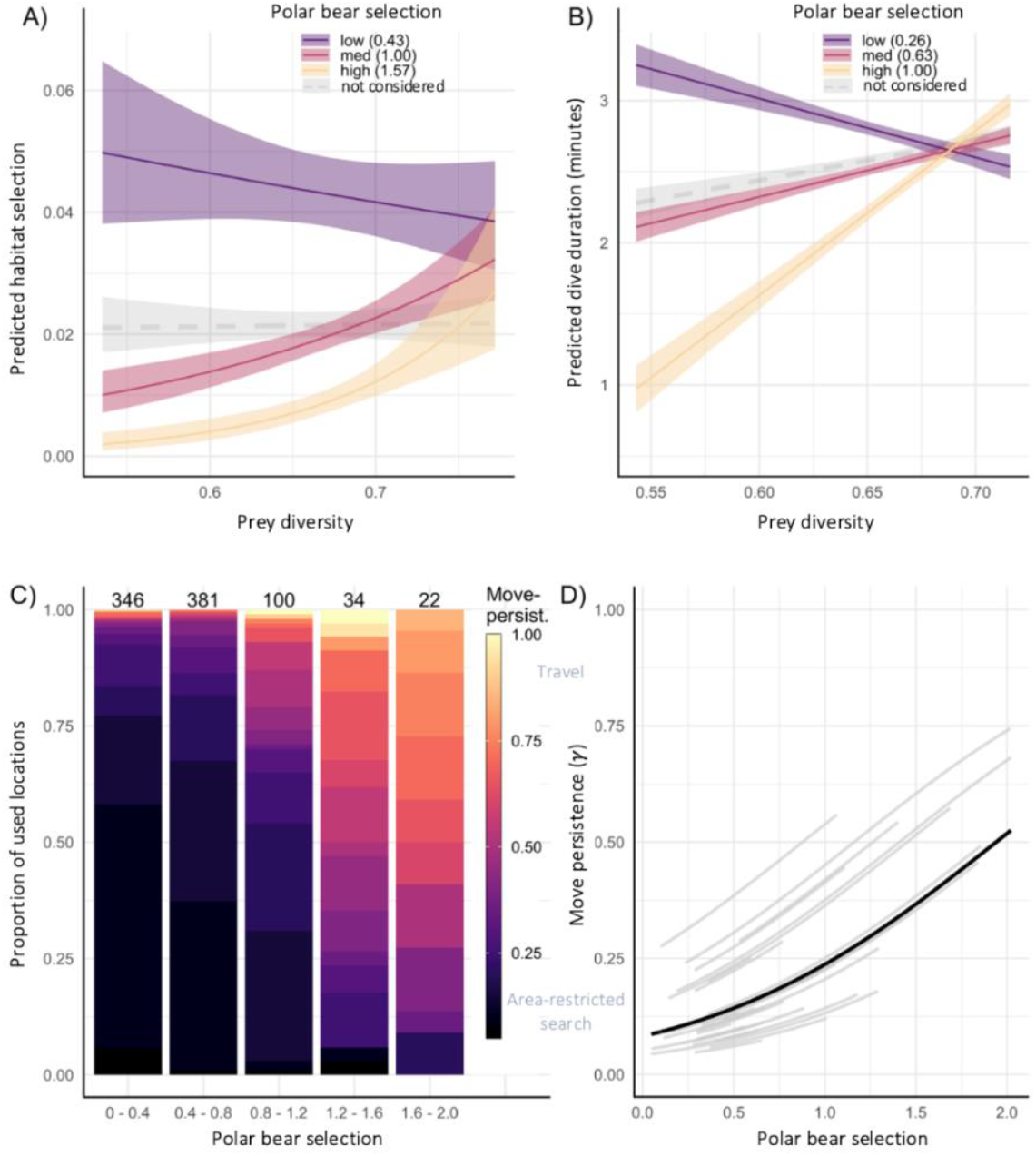
Predicted relationships between prey diversity, ringed seals, and polar bears. Predicted ringed seal (**A**) habitat selection from the best resource selection function model and (**B**) dive duration as a function of the interaction between polar bear selection and prey diversity, where the other covariates are held constant at their mean. Grey dashed line represents the model predictions where polar bear is excluded as a term in the model to highlight the insight that incorporating polar bear selection provides; where the top model was improved by 146 ΔAIC and 23 ΔAIC in (A) and (B), respectively (see Table S4). Shaded areas represent the 95% confidence intervals. (**C**) Proportion of estimated ringed seal behaviors (move-persistence values) relative to predation risk (polar bear selection). Move persistence values closer to zero (dark purple) are indicative of residency behaviors (e.g., area-restricted search behaviors such as foraging) and values closer to one (yellow) are indicative of travel. Polar bear selection values are split into equal sized groups where the groups are delineated on the x-axis. Sample size is presented for each polar bear selection group at the top of the bar. (**D**) Results from the ringed seal move persistence mixed model for fixed (black line) and random (individual seals, gray lines) effects, which shows that as predation risk (polar bear selection) increases, so does ringed seal move persistence (more persistent behaviors are likely more travelling-like behavior).

**Fig. 4.**
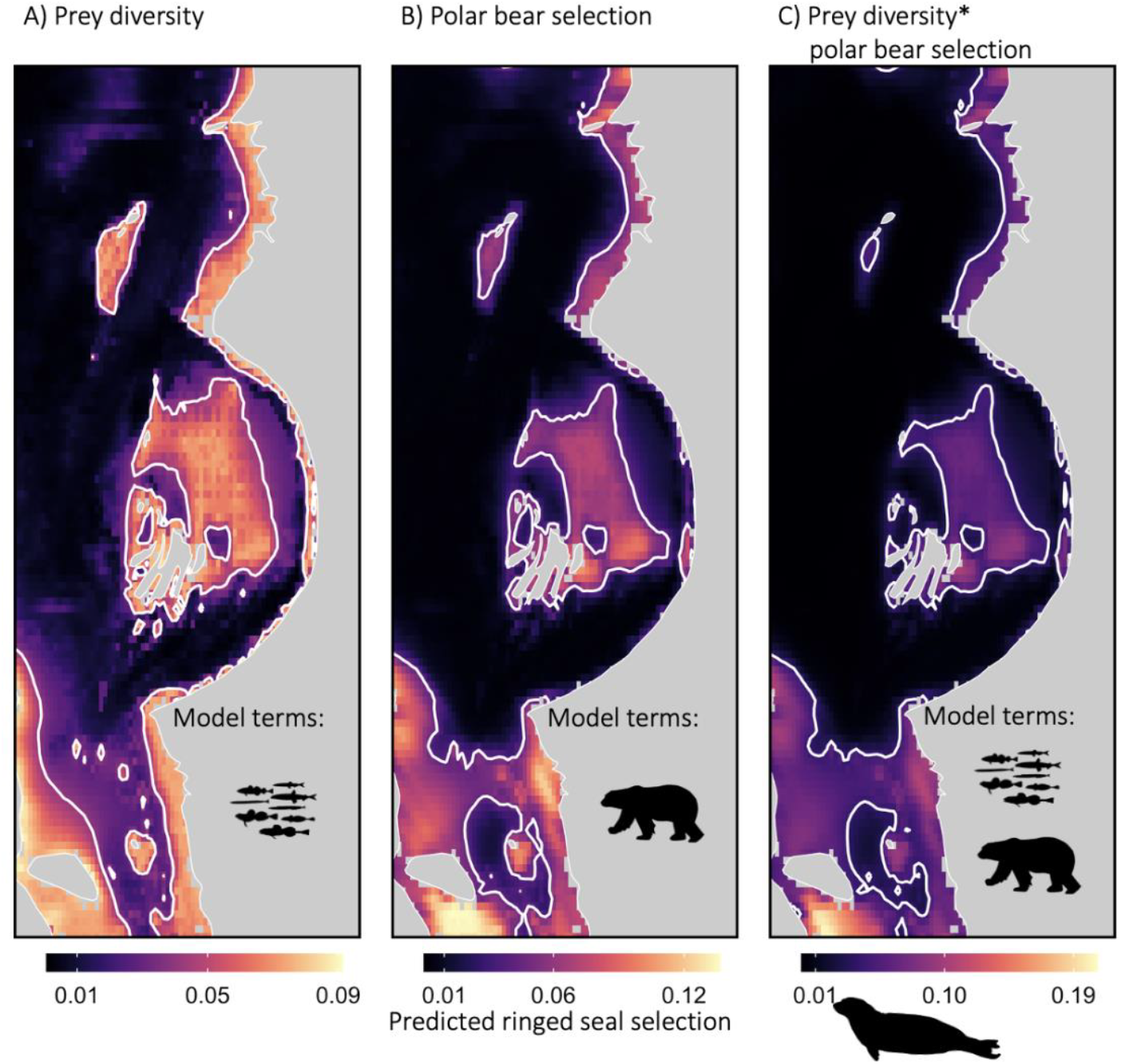
Predicted ringed seal habitat selection from various models. Predicted ringed seal selection from models that included covariates for (**A**) only prey diversity, (**B**) only polar bear selection, or (**C**) the interaction between prey diversity and polar bear selection (our best-supported model). Each model also included sea ice concentration and bathymetry as covariates. Covariates used to make predictions were taken for an example day, 15 February 2013. White contour lines delineate the 75% quantiles of predicted ringed seal selection for each panel.

Polar bear selection was a significant predictor of ringed seal movement behavior. We found that ringed seals spent more time in residency behaviors (i.e., low move-persistence, area-restricted search) when in areas with low polar bear selection values, and more time traveling when in areas of high polar bear selection values (Fig. 3C,D). Additionally, area-restricted search behaviors were virtually absent when in areas with high polar bear selection values (Fig. 3C).

### Ringed seal diving behavior

Our best-supported model for ringed seal dive duration included an interaction between polar bear selection and prey diversity (Table S4). This model suggested that, as polar bear selection increased, the duration of dives was more positively related to prey diversity (Fig. 3B). That is, when polar bear selection was low, dive duration decreased with increasing prey diversity. However, under conditions of moderate polar bear selection, dive duration increased with prey diversity. There was an even more drastic increase in dive duration with prey diversity when polar bear selection was high (Fig. 3B). This model also included sea ice concentration and dive depth as significant covariates, where dives were shorter in areas of high sea ice concentration and longer when dives were deeper (Table S5).

We did not find the expected negative relationships between polar bear selection and ringed seal dive frequency or ascent speed; rather the best-supported models for these dive metrics did not include the polar bear covariate (Table S4). The best-supported model for dive frequency included positive estimates for prey diversity and dive depth and a negative estimate for sea ice concentration, and the best-supported model for dive ascent speed included a positive estimate for dive depth (Table S5).

## Discussion

Our study explores the interactions between top-down and bottom-up processes that shape animal behavior and highlights the tradeoffs made by mesopredators under intense pressure from a specialized predator. Our results suggest that polar bears affect their primary prey, ringed seals, beyond consumptive effects by creating a landscape of fear that affects their habitat selection, movement ecology, and diving behavior. These results highlight the tradeoffs between predation risk and foraging, where ringed seals are more likely to be present and dive longer in high-risk areas when prey diversity is high. This interaction suggests a conditional response: seals generally avoid risky areas, but increasingly tolerate risk when prey diversity is high – likely reflecting greater tolerance in exchange for higher prey availability and accessibility, and thus foraging gains. Further, accounting for predators is a necessary consideration when examining the spatial ecology and behavior of prey. More broadly, our results exemplify how overlooking top-down pressures can bias ecological inference and potentially misguide conservation efforts in other predator-prey systems.

As predicted, and supported by theory (1, 62), the interaction between the prey and polar bear covariates suggests a higher selection by seals in areas with low predation risk and high-prey diversity. We also found that ringed seals were up to 5x more likely to use areas with low polar bear risk when prey diversity was low compared to when risk was high, and they appear to select riskier areas when prey diversity was high, conforming with the hazardous duty pay hypothesis which suggests that prey balance fear and food and that areas that have diverse resources afford sufficient fitness benefits or energetic returns to offset increased predation risk (6). In areas of low predation risk (i.e., low polar bear selection values), we found dive duration was negatively related to prey diversity, but in areas of high risk, dive duration was positively related to prey diversity (Fig. 3). More generally, we found that ringed seals dove for a shorter duration when in high-risk areas (i.e., ∼ 1 min in high-risk areas compared to ∼3 min in low-risk areas; unless prey diversity is high, Fig. 3B), perhaps to allow for time to return to an alternative breathing hole or haul-out lair if they detected a polar bear. If we consider dive duration as a proxy for time spent foraging (e.g., (63)), our results may suggest implications for energetic gain imposed by perceived predation risk (e.g., (64)). Taken together, our results suggest that ringed seals may tolerate high-risk areas when prey diversity exceeds a threshold (∼0.68, Fig. 3B), likely reflecting elevated foraging rewards. Prey diversity likely functions both as a biologically meaningful driver of seal behavior and as a proxy for prey availability across seasons. For generalist foragers like ringed seals, whose prey communities shift seasonally (65, 66), diversity may offer a more robust and ecologically relevant signal than single-species biomass. This dual role is supported by studies showing that prey diversity enhances predator biomass and feeding efficiency (33, 34, 67), and stabilizes resource availability across time (31, 32).

Our movement models and activity budgets suggest that low move-persistent behaviors conclude or are avoided when risk is high. Low move-persistence may reflect area-restricted search and is often associated with foraging; however, it may also represent a composite of resident movement patterns including resting, sleeping, intraspecific interactions, or predator avoidance (e.g., (30, 68, 69)). Regardless of the specific behavior, low move-persistence is associated with residency, and our results show it to be explained by our proxy of predation risk. The reduction of area-restricted search and increase of directed movements (i.e., traveling) associated with increasing polar bear selection values suggests that animals reduce these residency behaviors and instead travel through areas of high risk. Thus, the top-down pressure appears to shape the activity budgets of ringed seals. These results further concur with our findings of tradeoffs in diving where polar bears are associated with reduced time spent foraging (i.e., dive duration). Such behavioral responses to predation risk likely have an overall net positive effect on ringed seal survival, since they may reduce the risk of being killed by a polar bear. However, these risk-mitigation strategies likely come at a cost, and the demographic consequences of non-consumptive effects are unknown (70). Our results suggest that ringed seals may travel further to reach low-risk or high-reward sites and/or may forego foraging in areas that may have good, but not great, foraging payoff but have high risk. Any of these possibilities would reduce the average energy intake and increase the average energy expense of seals, leading to potential downstream effects on survival and reproduction (6). In extreme situations, ringed seals may become sufficiently undernourished to make riskier decisions, thus increasing mortality due to predation (71, 72).

Our study included both prey and predator modeled data in our ringed seal habitat selection, movement, and diving models, and the inclusion – or omission – of our proxy of predation risk altered ecological inference about ringed seals-prey relationships. For example, if polar bear selection was not included as an interacting term with prey diversity in our habitat selection or dive duration models, prey diversity would be interpreted as an unimportant variable (grey dashed line in Fig. 3A, B, and see (30)). Once polar bear selection was included as a covariate, the model revealed a possible predator-mediated relationship between ringed seals and their prey, demonstrating the importance of including top-down pressures when studying the ecological response of species to their prey and environment. This interaction reinforces our interpretation of prey diversity as both a biologically meaningful predictor and a proxy for prey availability, whose relevance emerges only when predation risk is accounted for.

Prediction maps can serve as an invaluable resource for managers to identify areas of conservation priority (e.g., protected areas, (12, 73). If models do not adequately represent species’ ecology, including the interactions between top-down and bottom-up processes, the subsequent maps can be less effective. Excluding the polar bear covariate from the model resulted in prediction maps that overestimated the size of areas characterized as most important for ringed seals and delineated different locations of the study region (Fig. 4A). Specifically, the prediction map based on the model that ignored predator risk (Fig. 4A) overestimated ringed seal selection in the north-central regions of the study area, and underestimated areas in the southern region (James Bay; relative to Fig. 4B-C). Given the extensive socio-economic challenges associated with creating protected areas and applying management policies (74), it is critical to provide the most accurate information (13) and thus incorporate these important dynamics.

Under widespread climate change and habitat alteration, insight into complex – and likely changing – trophic interactions is important for understanding how species’ ecology may shift. For example, many species globally are suffering from reduced foraging success (75) and under the starvation-predation hypothesis, animals in poor body condition may forego anti-predator behaviors to focus on foraging (71, 72, 76). In other cases, the influx of new (or more common) predators may provide additional predation pressures (22, 77). The changing habitat and anthropogenic pressures may alter predator-prey relationships and movement (8); for example, the increase in roads and seismic lines has intensified gray wolf (*Canis lupus*) predation on boreal caribou (*Rangifer tarandus caribou*, (78)). Further, large carnivores have undergone population declines globally (79), resulting in weakened landscapes of fear (80). Humans can also shape landscapes of fear, either increasing perceived risk through disturbance, or in some contexts, reducing predation risk for prey when predators avoid human activity (i.e., “human shield hypothesis”, (81)). In our system, both ringed seals and polar bears are vulnerable to climate-mediated reductions in sea-ice habitat (23, 24). The projected reductions in polar bear abundance may weaken the landscape of fear (e.g., as in the case of reduced Tasmanian Devils, *Sarcophilus harrisii*, (82)) and the expected increase in prey diversity (29) may increase foraging opportunities for seals. However, these benefits for ringed seal populations will likely be offset by the demographic consequences of reduced sea ice (25), and the predicted smaller body size and lower energetic value of prey (29). Polar bears have evolved as a specialized predator, and in response, ringed seals have evolved antipredator behaviors. Our observations of polar bear habitat selection explaining ringed seal habitat selection and dive duration may suggest that ringed seals have tactics of identifying high-risk areas. As reduced sea ice results in decreased polar bear population sizes, and increased access for other predators, such as killer whales (83), the usefulness of these antipredator tactics by seals is unknown. Further, reductions in the size of the foraging arena (i.e., sea ice extent) may inflate relative polar bear density in the interim, increasing top-down pressures (84). Thus, our observations of reduced foraging behavior in high-risk areas may contribute to observed reductions in ringed seal body condition (25). Future work should assess whether prey diversity continues to buffer risk under shifting predator regimes and declining habitat quality, especially given projections of increased prey diversity but reduced fish body size by the end of the century (29).

We expect ringed seals to avoid polar bears, but we also would expect polar bears to match ringed seal distribution (i.e., a positive relationship) through predator-prey evolution. We suggest that while ringed seals are the primary food source for polar bears – especially females throughout the fall, winter, and spring– they are not the only source (e.g., (17)) and other seasonal prey distributions, as well as habitat requirements (e.g., adequate ice for travel and foraging) also likely influence the habitat use of polar bears across their large home ranges (85, 86). In other systems, low-risk areas are associated with areas that are not conducive to prey capture (e.g., (87, 88)). In our study, ringed seals may employ predator-avoidance strategies in the early winter (mid-December through mid-January) by staying in open water, and then shift to antipredator behaviors – such as modifying their diving – throughout the winter when Hudson Bay freezes over, low-risk areas shrink, and open water is not available (Fig. S4, S5). Open water generally limits polar bear’s movement and ability to successfully hunt seals, which may explain the seal selection for open water in the early winter (85, 89). Polar bears and ringed seals may play a dynamic game of cat and mouse where the “behavioral race” is dependent on the mobility and response of both the predator and prey (90); future work should explore the potential lag effects between predation risk and mesopredator habitat selection, especially under shifting ice conditions. Integrating mapped kill sites, aerial survey data of breathing holes, and polar bear tracks could help disentangle perceived risk, behavioral avoidance, and actual predation events.

Our models performed well according to our visual assessment of residuals and cross-validation exercise (Figs. S1-S3, Table S6-S8), but they are inherently subject to limitations. Argos data are associated with large location error (35). Thus, we applied a state-space model (36) to the Argos data, which is an approach that has been shown to improve the location quality (91), and propagated the error in the final resource selection function model (Table S6). However, newer tags (i.e., Fastloc-GPS, (91)) can provide more accurate locations at a finer temporal resolution that could be used to investigate smaller-scale patterns in movement and diving and remove much of the uncertainty associated with Argos data. Additionally, our prey and polar bear covariates were modeled predictions that are unlikely to be perfectly matched the distribution of these species, and the prey data were modelled at a relatively coarse spatial resolution. However, both approaches have performed well in validation exercises (29, 30, 39), the prey datasets have significantly outperformed other fine-scale proxies of prey abundance (30), and prey diversity was consistently included in our highest-ranking models (Table S4). We acknowledge that our proxy polar bear predation risk was calculated using sea ice concentration and bathymetry as covariates, and that our models also included these covariates, but none of the models’ covariates were strongly correlated. We did not find the hypothesized relationships between polar bear selection and ringed seal dive frequency or ascent speed; perhaps ringed seals can only detect cues of polar bear presence within the last few meters of ascent, and future work using more fine scale biologging tools (e.g., videologgers) could investigate dives at a resolution that would allow identification of surfacing behavior. Additionally, the use of accelerometers that can detect changes in buoyancy – an indication of body condition – during passive drift dives could offer further insight into the energetic consequences of predator avoidance. Finally, the polar bear layers were generated using movement data from only female polar bears due to limitations in placing a collar on a male bear (their neck is wider than their head). Although there are known differences in movement patterns between male and female polar bears, they have largely similar movement patterns (92) and sea ice habitat selection (93). Further, ringed seals generally contribute more to the diet composition of female than male polar bears (17) and thus female polar bears might impose more of a predation risk.

Predator avoidance is observed across many taxa (e.g., barnacle larvae *Semibalanus balanoides*: (94), northern elephant seals *Mirounga angustirostris:* (95), ibex *Capra sibirica* and argali *Ovis ammon* (96)) and how species navigate the critical trade-off between predation risk and resource acquisition is a fundamental question in ecology, however, combined top-down and bottom-up processes are seldom incorporated into the same models. Our findings fill knowledge gaps on the complex dynamics between predators and prey and advance our understanding of non-consumptive effects of predation, providing empirical evidence of how species balance risk-reward trade-offs in extreme environments. Similar processes are likely occurring in other ecosystems where prey species must balance the trade-off between accessing resources and minimizing predation risk. For instance, species experiencing analogous selective pressures – such as migratory ungulates navigating risky landscapes (78) or marsupials modifying their foraging patterns to mitigate predation risk (82) – may exhibit comparable habitat selection patterns. Simultaneously considering the influence of predator and prey is important for ecological inference and thus should be considered when modeling habitat use and selection, especially for critical habitat designation. In a time of rapid climate change, habitat degradation, and anthropogenic disturbance, properly directing conservation efforts by explicitly accounting for both top-down and bottom-up processes in habitat selection models for better understanding of animal ecology is of utmost importance.

## Acknowledgments

We gratefully acknowledge the Inuit hunters and the Sanikiluaq Hunters and Trappers Association for assistance in the field, especially Lucassie and Johnassie Ippak and their families. We thank Brian Hunt, Chris Harley, David Rosen, Aaron Fisk, two anonymous reviewers, and the Editor for their constructive and thoughtful feedback on earlier versions of this manuscript. Ringed seal capture and handling procedures were approved annually by the Freshwater Institute Animal Care Committee (FWI-ACC-2006 to 2012) and Fisheries and Oceans Canada’s Licenses to Fish for Scientific Purposes (S-05/06 - 09/13-1006-NU). Polar bear capture and handling procedures were approved annually by the Animal Care Committees of the Ontario Ministry of Natural Resources (OMNR ACC 2006-95 to ACC 2014-95) and York University (Protocol #s 2012-12W and 2014-5W).

## Funding

Fisheries and Oceans Canada (SF, DY)

Nunavut Wildlife Management Board (SF, DY)

ArcticNet (SF, DY)

Natural Sciences and Engineering Research Council of Canada (NSERC) (All authors)

Weston Family Foundation (KF)

Polar Knowledge Canada (KF)

Earth Rangers (KF)

Canada Research Chairs program (MAM)

Canada Foundation for Innovation (MAM)

B.C. Knowledge Development Fund (MAM)

OMNR’s Wildlife Research and Development Section (MO, GT, JN)

OMNR’s Climate Change Program (MO, GT, JN)

OMNR’s Species at Risk Research Fund for Ontario (MO, GT, JN)

Polar Bears International (MO, GT, JN)

Toronto Zoo’s Endangered Species Reserve Fund (MO, GT, JN)

Helen McCrea Peacock Foundation (MO, GT, JN)

Canadian Wildlife Federation (MO, GT, JN)

Born Free Foundation (MO, GT, JN)

Wildlife Conservation Society Canada (MO, GT, JN)

