## Appendix for "Top-down and bottom-up processes jointly explain mesopredator movement and foraging ecology"

### Supplementary Material

Supplementary Methods

*Prey biomass predictions*

The modelled prey biomass and diversity data were obtained from (1). Briefly, (1) used a dynamic bioclimate envelope model (DBEM) to predict the abundance, biomass (in tonnes), and distribution of prey species from ringe seals (*Pusa hispida*) in the Hudson Bay System from 1950 to 2100 based on changes in ocean conditions (e.g., sea temperature, pH, salinity). Unlike other methods that rely on functional groups (see for model comparisons: (2, 3)), the DBEM used species-specific physical and biological conditions to identify suitable habitats for eight main prey species of ringed seals: Arctic cod (*Boreogadis saida*), capelin (*Mallotus villosus*), Pacific sand lance (*Ammodytes hexaoterus*), Northern sand lance (*Ammodytes dubius*), rainbow smelt (*Osmerus mordax*), Arctic staghorn sculpin (*Gymnocanthus tricuspis*), shorthorn sculpin (*Myoxocephalus Scorpius*), and Moustache sculpin (*Triglops murrayi*). The DBEM used empirical species occurrence data and associated biogeographic characteristics to establish baseline data; since empirical data does not exist for the Hudson Bay System, baseline data were realized through empirical data from other regions, allowing the model to describe relative differences in abundance and biomass between species and through space (4). We predicted species abundance, biomass, and distribution under two different climate change scenarios: a low- and high-emission scenario. The low-emission scenario corresponds with strong-mitigation to limit global warming below 2°C by 2100, and the high-emission scenario corresponds with little to no mitigation to limit global warming (business as usual). Modelling these two emission scenarios allows for scaling the results to in-between scenarios, however, we used the high-emission scenario as it aligns closely to emissions during our study period (5, 6).

*Predicting polar bear predation risk*

We used data from 39 female polar bears radio-collared in southern Hudson Bay between 2007 and 2013. We chemically immobilized bears from a helicopter by remote drug delivery using either a combination of xylazine and zolazepam–tiletamine (XZT) or a combination of medetomidine and zolazepam–tiletamine (MZT). XZT was administered as xylazine (Cervizine 300®, Wildlife Pharmaceuticals, Inc., Fort Collins, Colorado, USA) at 2 mg/kg and Telazol® (Fort Dodge Laboratories, Inc., Fort Dodge, Iowa, USA) at 3 mg/kg estimated body mass (7). MZT was administered as medetomidine (Bow River Pharmaceuticals, Bow River, Alberta, Canada) at 0.06 mg/kg and Telazol® at 2 mg/kg (8). At the conclusion of handling, we administered atipamezole (Antisedan®, Pfizer Bio-Pharmaceuticals and Animal Health, Mississauga, Ontario, Canada) at 0.20 mg/kg to reverse xylazine or 0.30 mg/kg to reverse medetomidine (7, 8). Handling procedures followed the general guidelines of the Canadian Council for Animal Care (CCAC 2003) and the American Society of Mammalogists (9). Each year we deployed Telonics Gen III (TGW-3680) or Gen IV (TGW-4680, TGW-4670) GPS telemetry collars (Telonics Inc., Mesa, Arizona, USA) programmed to record a GPS position every 4 hours (i.e., six locations per day) on a sample of females either with cubs of the year or with yearlings. Data were delivered through the Argos Direct Automatic Distribution Service

(ADS) or Iridium network. All collars also stored locations in memory which could be downloaded upon recovery. Collars were fitted with timed release mechanisms (CR-2A; Telonics Inc, Mesa, Arizona), programmed to drop off after either 1 (Gen III) or 2 (Gen IV) years.

To predict a proxy of predation risk, we fit a resource selection function (RSF) to the polar bear data. We used bathymetry, daily sea ice concentration, and distance to coast as covariates to predict predation risk because analysis of their previously documented importance for estimating on-ice polar bear habitat selection (10–13). Bathymetry (m) was extracted from NOAA's ETOPO1 Global Relief Model (~1 km resolution and aggregated at a 5 km resolution). Daily sea ice concentration (0-100%) was provided from the Nimbus-7 SSMR and DMSP SSM/I-SSMIS Passive Microwave (25 x 25 km resolution). Distance to coast (m) was derived from the bathymetry centroids. Each environmental covariate was extracted for each polar bear location found within our study area and the ice season (from fall freeze-up to spring break-up, n = 18,226 locations). The availability sample was created by sampling each 5 km x 5 km cell within the study area (-82.0 to -76.0°W and 52.7 to 61.0°N). We first calculated mean bathymetry within each cell and placed a single point within the middle of the cell in order to calculate the distance of the cell's centroid to the nearest coastline. These same points were then used to extract daily sea ice concentration, as above, for each day of our study period. We also allowed the effects of each variable to change in response to changing sea ice concentration (i.e., functional response), which has been shown to offer more robust predictions of species distribution, particularly in highly dynamic environments such as Arctic sea ice (14–16). Thus, the covariates in our full model were as follows:

$$IceCon + IceCon^2 + Bath + Bath^2 + DistCoast + DistCoast^2 + IceConc \cdot MeanIceConc + IceConc^2 \cdot MeanIceConc + Bath \cdot MeanIceConc + Bath^2 \cdot MeanIceConc + DistCoast \cdot MeanIceConc + DistCoast^2 \cdot MeanIceConc,$$

where *MeanIceConc* represents the mean daily sea ice concentration for the entire study extent.

The final model was fitted using a Least Absolute Shrinkage and Selection Operator (LASSO) framework using the R package glmnet (17, 18). Specifically, the model was fitted using weighted logistic regression and 10-fold cross-validation to select the optimal value of lambda (the tuning parameter used to regularize coefficient values).

<u>Supplementary Results</u>

*RSF models and validation*

The best-supported prey-only model included prey diversity as a covariate (Table S2). Thus, prey diversity was selected as the prey covariate in all subsequent analyses. We did not find any large correlation coefficients between the polar bear, prey diversity, bathymetry, or sea ice concentration covariates and thus allowed them to be included together in candidate models (Table S3). We found 25 available points per one used location resulted in stabilized parameter estimates (Fig. S1).

Figures

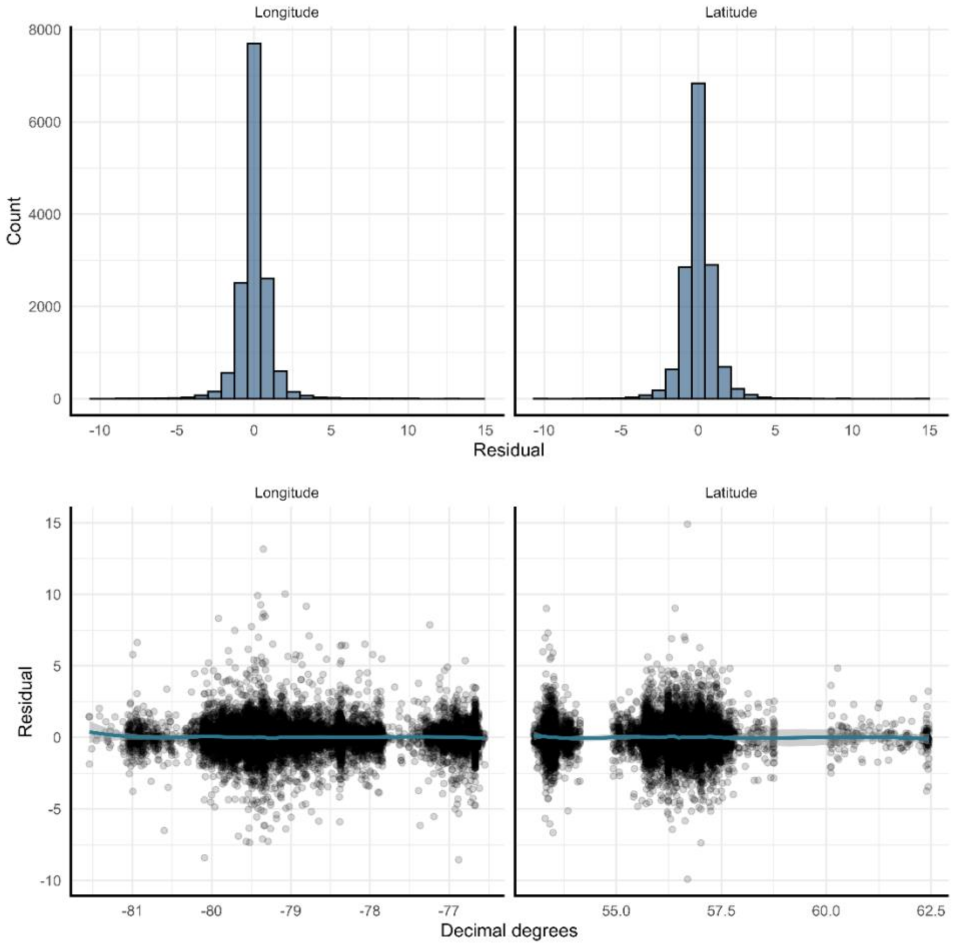


**Figure S1**. One-step-ahead residuals from the ringed seal state space model for both longitude (left) and latitude (right) presented as histograms (top) and plots as a function of decimal degrees (bottom).

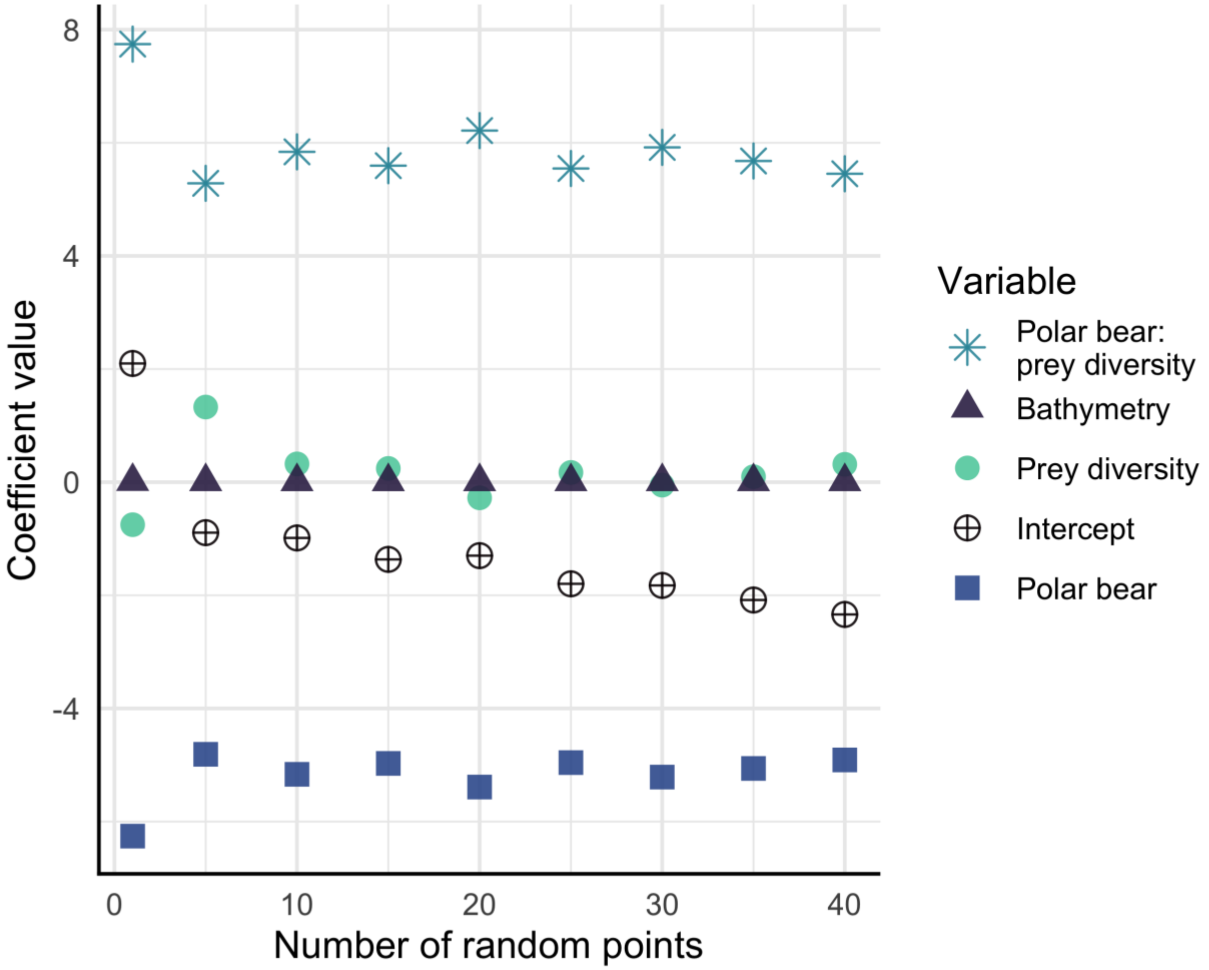


**Figure. S2**. Coefficient estimates for our best-supported RSF with various amounts of random points in the availability sample, which included bathymetry, polar bear selection, prey diversity, and the interaction between polar bear selection and prey diversity. The coefficients stabilize after ~25 random points (available locations) are generated per used point.

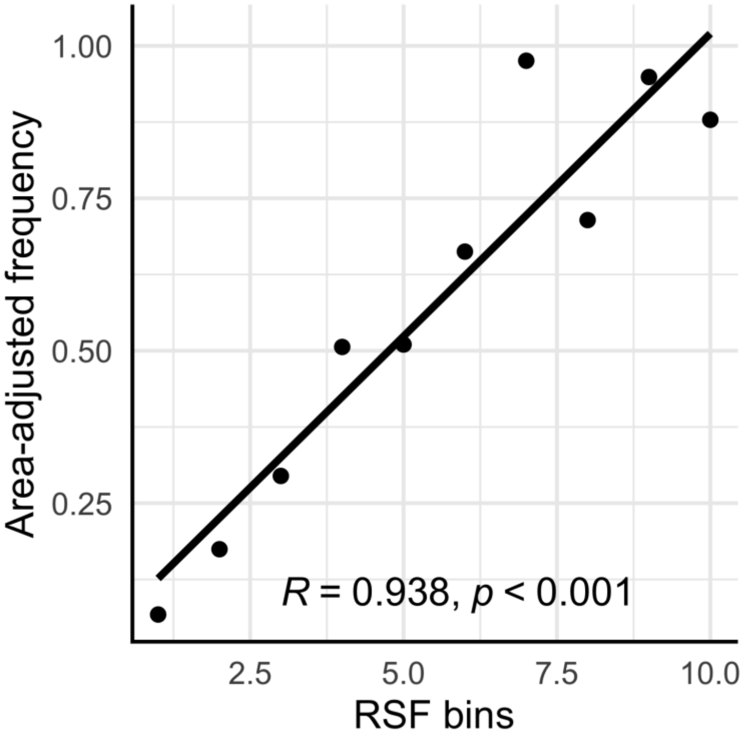


**Figure. S3**. Results from leave-one-individual-out cross validation on the best-supported RSF which included bathymetry and sea ice concentration as well as the interaction between polar bear selection and prey diversity. Spearman Rank correlation coefficient is presented on the plot. RSF bins represent the 10 even-increments of predicted ringed seal selection (e.g., 1-2, 2-3, 3-4, etc.). The black line represents the line of best fit.

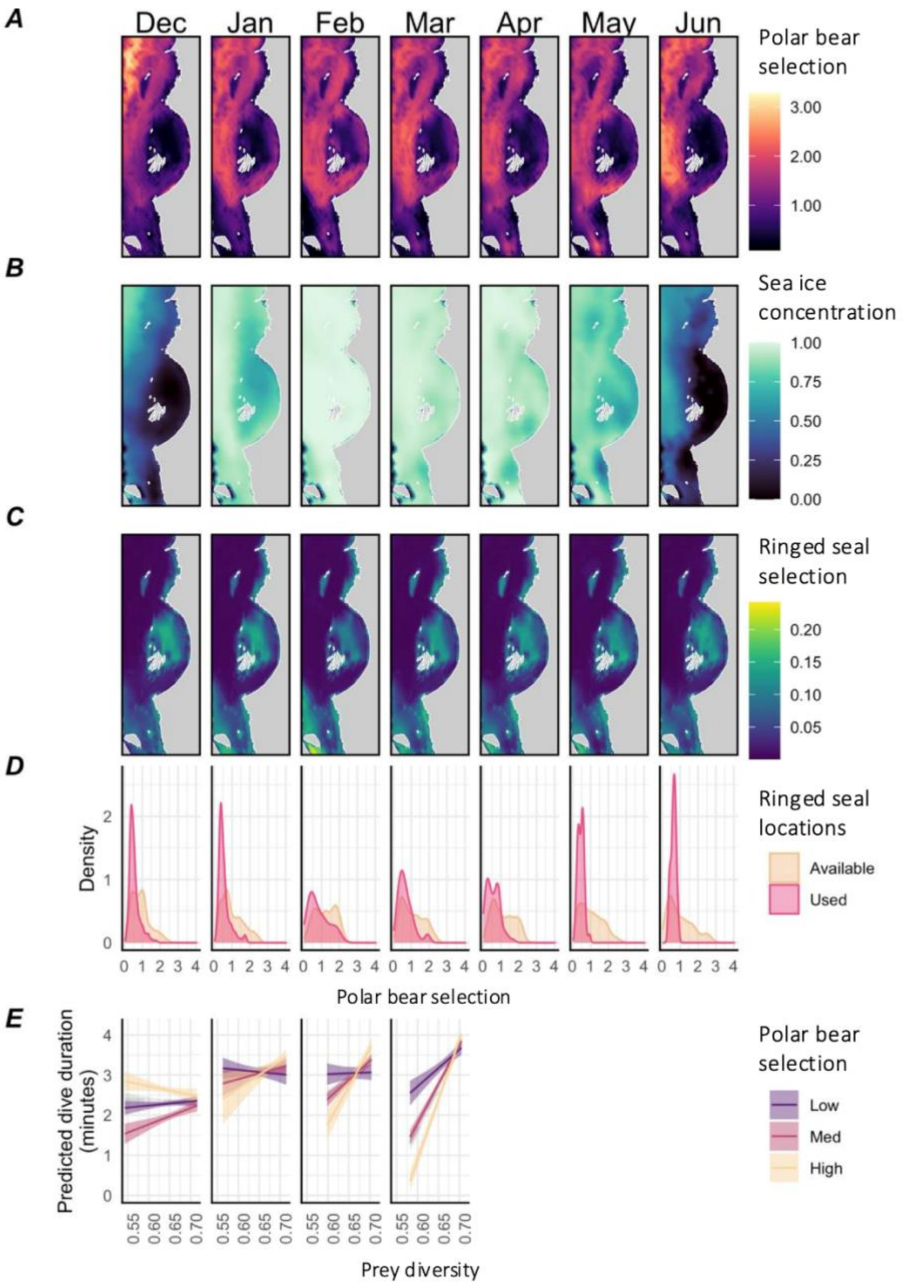


**Figure. S4**. A) Predicted polar bear selection, B) sea ice concentration, C) predicted ringed seal selection from the best model (which included an interaction between prey diversity and polar bear selection, as well as sea ice concentration and bathymetry as covariates), D) Density of ringed seal used or available locations used in resource selection functions as a function of polar bear selection in those locations, E) dive duration as a function of the interaction between polar bear selection and prey diversity, where the other covariates are held constant at their mean, where the shaded areas represent the 95% confidence intervals; note that small samples sizes in Apr-Jun precluded month-specific model fitting. We selected the 15th of each month (averaged amongst years) for A-C, and all monthly data were pooled for D-E.

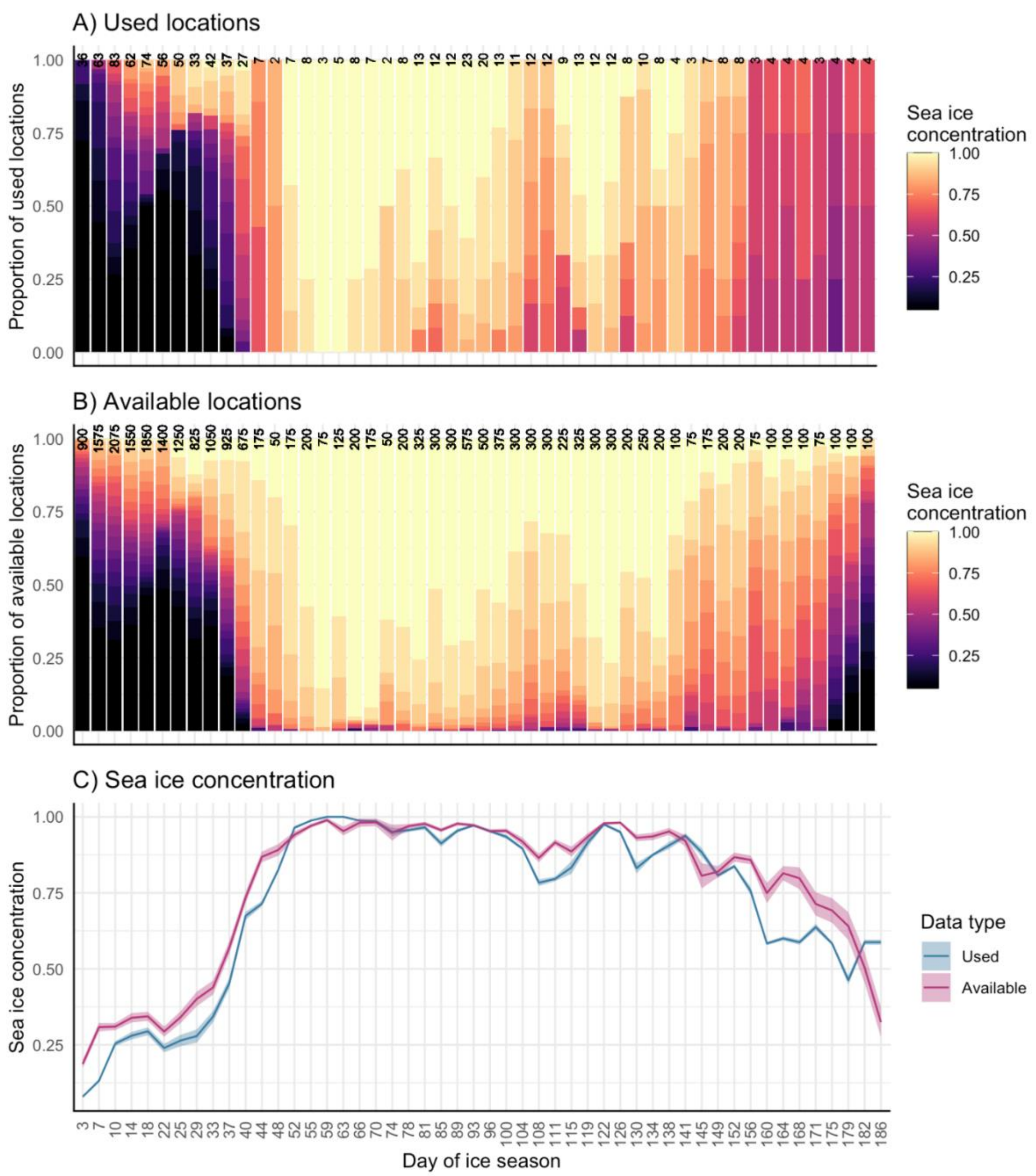


**Figure S5**. Proportion of A) used or B) available locations of ringed seals (1 or 25 per seal per day, respectively) relative to day of ice season, grouped by sea ice concentration. C) Sea ice concentration (mean +/- 95% confidence intervals) for used and available locations. These data were used in the resource selection functions. Day of ice season starts represents days since freeze-up (approx. mid-December, see *Predicting polar bear selection*). Day of ice season is split into equal sized groups where the groups of day of ice season are delineated on the x-axis and sample size is presented for each group at the top of the bar in A and B.

**Table S1**. Summary of ringed seal movement and dive data used in analysis. Note that the location counts represent the numbers predicted by the state-space model. Filtered dives (n) were spatially and temporally interpolated to predicted locations.

| **ID** | **Sex** | **Start date** | **End date** | **Duration** | **Predicted locations (n)** | **Filtered dives (n)** |
|---|---|---|---|---|---|---|
| 106373 | male | 2011-12-10 | 2012-03-17 | 99.0 days | 505 | 5817 |
| 106384 | male | 2011-12-10 | 2011-12-21 | 11.0 days | 134 | 421 |
| 106385 | female | 2011-12-10 | 2012-02-02 | 54.0 days | 262 | 2868 |
| 106387 | male | 2011-12-10 | 2011-12-19 | 8.8 days | 107 | 478 |
| 106388 | male | 2011-12-10 | 2011-12-23 | 13.0 days | 157 | 812 |
| 107829 | female | 2011-12-10 | 2012-03-18 | 100.0 days | 295 | 1416 |
| 116482 | female | 2012-12-07 | 2012-12-19 | 13.0 days | 156 | 1244 |
| 116483 | male | 2012-12-11 | 2013-04-02 | 110.0 days | 657 | 7510 |
| 116484 | male | 2012-12-07 | 2013-04-19 | 130.0 days | 715 | 4153 |
| 116486 | male | 2012-12-07 | 2013-05-14 | 160.0 days | 464 | 3388 |
| 116488 | female | 2012-12-12 | 2012-12-22 | 10.0 days | 122 | 679 |
| 116489 | male | 2012-12-07 | 2013-01-16 | 40.0 days | 478 | 3810 |
| 116491 | female | 2012-12-07 | 2012-12-27 | 20.0 days | 240 | 3385 |
| 116493 | male | 2012-12-16 | 2012-12-28 | 12.0 days | 142 | 947 |
| 116495 | female | 2012-12-07 | 2013-06-13 | 190.0 days | 1726 | 14844 |
| 116496 | female | 2012-12-11 | 2012-12-24 | 13.0 days | 160 | 834 |
| 43835 | male | 2010-12-15 | 2011-01-18 | 35.0 days | 357 | 2253 |
| 43836 | female | 2010-12-21 | 2011-01-17 | 27.0 days | 330 | 1385 |
| 43838 | female | 2010-12-23 | 2011-01-14 | 22.0 days | 249 | 1204 |
| 43842 | female | 2010-12-15 | 2011-01-17 | 34.0 days | 403 | 1952 |
| 43845 | male | 2010-12-15 | 2011-04-25 | 130.0 days | 818 | 4446 |
| 43852 | female | 2010-12-15 | 2011-01-18 | 34.0 days | 406 | 1932 |
| 43853 | female | 2010-12-20 | 2011-01-11 | 21.0 days | 257 | 1165 |
| 43857 | female | 2010-12-15 | 2011-02-26 | 73.0 days | 514 | 3755 |
| 43858 | female | 2010-12-15 | 2010-12-31 | 17.0 days | 203 | 603 |
| 43865 | female | 2010-12-29 | 2011-01-12 | 13.0 days | 162 | 641 |
| **TOTAL** | | | | **1389.8 days** | **10019** | **71942** |

**Table S2**. Results from resource selection function models for ringed seal movement locations in southeastern Hudson Bay. Each model had a different prey metric as the fixed effect. Models were ranked using AICc to determine which prey metric best explained used locations; the best prey metric was used in subsequent modeling. Each prey metric was scaled prior to modeling.

| **Explanatory variable** | **AICc** | **ΔAICc** | **Est** | **SE** | ***p*-value** |
|---|---|---|---|---|---|
| Prey diversity | 7656.72 | 0.00 | 0.402 | 0.038 | <0.001 |
| Biomass of all prey species | 7751.54 | 94.82 | 0.174 | 0.035 | <0.001 |
| Biomass of capelin | 7776.36 | 119.64 | -0.027 | 0.034 | 0.423 |
| Biomass of Arctic cod | 7776.52 | 119.80 | -0.023 | 0.034 | 0.488 |
| Biomass of northern sand lance | 7776.98 | 120.26 | -0.004 | 0.034 | 0.888 |

Est = parameter estimate, SE = standard error of parameter estimate

**Table S3.** Correlation coefficients between covariates for A) movement data that was used for the resources selection function (RSF), and B) diving data that was used for the linear mixed effects models (LMEs).

| **A) Movement data for RSF** | | | | |
|---|---|---|---|---|
| | **Polar bear** | **Prey diversity** | **Ice concentration** | **Bathymetry** |
| **Polar bear** | 1 | | | |
| **Prey diversity** | -0.22 | 1 | | |
| **Ice concentration** | 0.45 | 0.00 | 1 | |
| **Bathymetry** | -0.59 | 0.43 | -0.10 | 1 |
| **B) Diving data for LMEs** | | | | |
| | **Polar bear** | **Prey diversity** | **Ice concentration** | **Dive depth** |
| **Polar bear** | 1 | | | |
| **Prey diversity** | 0.19 | 1 | | |
| **Ice concentration** | 0.22 | 0.28 | 1 | |
| **Dive depth** | 0.27 | -0.02 | 0.12 | 1 |

**Table S4**. Candidate models of ringed seal A) habitat selection, and dive B) frequency, C) ascent speed, and D) duration. Models were ranked by Akaike information criterion with small sample correction (AICc) where the "best" model (bolded) is that with the fewest number of parameters estimated and within two ΔAICc of the top model. We present candidate models when the AICc is less than that of the null model.

| **Terms** | **AICc** | **ΔAICc** |
|---|---|---|
| **A) Resource selection function** | | |
| ice concentration + polar bear*prey diversity + bathymetry | 7009.30 | 0.00 |
| **polar bear*prey diversity + bathymetry** | **7009.87** | **0.57** |
| polar bear + prey diversity + bathymetry | 7017.52 | 8.22 |
| ice concentration + polar bear + prey diversity + bathymetry | 7018.22 | 8.92 |
| polar bear + ice concentration + bathymetry | 7037.43 | 28.12 |
| polar bear + bathymetry | 7038.55 | 29.25 |
| prey diversity + ice concentration + bathymetry | 7155.30 | 146.00 |
| prey diversity + bathymetry | 7161.85 | 152.55 |
| ice concentration + bathymetry | 7166.25 | 156.95 |
| bathymetry | 7170.45 | 161.15 |
| **B) Dive frequency** | | |
| **prey diversity + ice concentration + depth** | **10644.8** | **0.0** |
| polar bear + prey diversity + ice concentration + depth | 10645.9 | 1.2 |
| ice concentration + polar bear*prey diversity + depth | 10647.1 | 2.3 |
| ice concentration + depth | 10647.9 | 3.2 |
| polar bear + ice concentration + depth | 10648.3 | 3.5 |
| prey diversity + depth | 10658.1 | 13.3 |
| depth | 10659.1 | 14.3 |
| **C) Dive ascent speed** | | |
| prey diversity + depth | 3187.5 | 0.0 |
| **depth** | **3189.2** | **1.7** |
| **D) Dive duration** | | |
| **ice concentration + polar bear*prey diversity + depth** | 84616.9 | 0.0 |

| | | |
|---|---|---|
| polar bear *prey diversity + depth | 84625.5 | 8.6 |
| polar bear + ice concentration + prey diversity + depth | 84628.9 | 12.0 |
| polar bear + prey diversity + depth | 84638.3 | 21.3 |
| polar bear + ice concentration + depth | 84639.5 | 22.6 |
| ice concentration + prey diversity + depth | 84639.9 | 22.9 |
| polar bear + depth | 84648.1 | 31.2 |
| ice concentration + depth | 84649.8 | 32.9 |
| prey diversity + depth | 84651.2 | 34.3 |
| depth | 84660.7 | 43.7 |

**Table S5**. Results from the best models (bolded in Table S6) for ringed seal A) habitat selection, and dive: B) frequency, C) ascent speed, and D) duration.

| | | | |
|---|---|---|---|
| **A) Resource selection function** | | | |
| polar bear | -5.120 | 1.002 | <0.001 |
| preydiv | 0.272 | 1.146 | 0.813 |
| polar bear: prey diversity | 5.964 | 1.516 | <0.001 |
| **B) Dive frequency** | | | |
| (Intercept) | 2.39 | 0.80 | 0.003 |
| ice concentration | -0.42 | 0.11 | <0.001 |
| prey diversity | 2.84 | 1.24 | 0.022 |
| **C) Dive ascent speed** | | | |
| (Intercept) | 0.60 | 0.02 | <0.001 |
| **D) Dive duration** | | | |
| (Intercept) | 135.53 | 104.38 | 0.194 |
| ice concentration | -19.64 | 9.19 | 0.033 |
| polar bear | -171.93 | 109.34 | 0.116 |
| prey diversity | -35.11 | 160.48 | 0.827 |
| polar bear: prey diversity | 230.95 | 162.56 | 0.155 |
| depth | 3.64 | 0.042 | <0.001 |

Est = parameter estimate, SE = standard error of parameter estimate

**Table S6**. Results from the best resource selection function for ringed seals accounting for uncertainty of the predicted locations. We ran posterior simulations for each seal track, fit the resource selection function, and repeated this process 100 times. We present the mean, 2.5$^{th}$, and 97.5$^{th}$ percentile of the parameter estimate.

| | **Parameter Estimate** | | |
|---|---|---|---|
| | **Mean** | **2.5$^{th}$ percentile** | **97.5$^{th}$ percentile** |
| polar bear | -5.13 | -5.30 | -4.98 |
| preydiv | 0.258 | 0.105 | 0.391 |

| | | | |
|---|---|---|---|
| polar bear: prey diversity | 5.98 | 5.75 | 6.23 |
| bathymetry | 0.011 | 0.011 | 0.012 |

**Table S7**. Resource selection function candidate models for ringed seals, using the dataset of all predicted ringed seal locations (i.e., at a 2-hr timestep). Models were ranked by Akaike information criterion with small sample correction (AICc) where the "best" model (bolded) is that with the fewest number of parameters estimated and within two ΔAICc of the top model. We present candidate models when the AICc is less than that of the null model.

| **Terms** | **AICc** | **ΔAICc** |
|---|---|---|
| **ice concentration + polar bear * prey diversity + bathymetry** | **78926.44** | **0.00** |
| polar bear * prey diversity + bathymetry | 78972.62 | 46.18 |
| polar bear + prey diversity + ice concentration + bathymetry | 79218.15 | 291.71 |
| polar bear + prey diversity + bathymetry | 79236.22 | 309.78 |
| polar bear + ice concentration + bathymetry | 79271.16 | 344.72 |
| polar bear + bathymetry | 79298.50 | 372.05 |
| prey diversity + ice concentration + bathymetry | 80734.87 | 1808.42 |
| ice concentration + bathymetry | 80739.84 | 1813.40 |
| prey diversity + bathymetry | 80813.65 | 1887.21 |
| bathymetry | 80814.12 | 1887.68 |

**Table S8**. Results from the best resource selection function model (bolded in Table S6) for ringed seals using the dataset of all predicted ringed seal locations (i.e., at a 2-hr timestep).

| | **Est** | **SE** | **p-value** |
|---|---|---|---|
| ice concentration | 0.198 | 0.028 | <0.001 |
| polar bear | -8.903 | 0.460 | <0.001 |
| prey diversity | -5.567 | 0.491 | <0.001 |
| polar bear : prey diversity | 11.149 | 0.671 | <0.001 |
| bathymetry | 0.011 | 0.001 | <0.001 |

Est = parameter estimate, SE = standard error of parameter estimate

**Table S9**. Candidate models of ringed seal move-persistence. Models were ranked by Akaike information criterion with small sample correction (AICc) where the "best" model (bolded) is that with the fewest number of parameters estimated and within two ΔAICc of the top model.

| **Terms** | **AICc** | **ΔAICc** |
|---|---|---|
| polar bear + prey diversity + bathymetry | -2240.3 | 0 |
| **polar bear + bathymetry** | **-2238.7** | **1.5** |
| prey diversity + bathymetry | -2236.5 | 3.7 |

**Table S10**. Results from the best-supported move-persistence mixed model (bolded in Table S8).

| | **Est** | **SE** | **p-value** |
|---|---|---|---|
| (Intercept) | -3.107 | 0.675 | <0.001 |
| Polar bear | 1.685 | 0.687 | 0.014 |
| Bathymetry | -0.001 | 0.007 | 0.956 |

Est = parameter estimate, SE = standard error of parameter estimate
